# Cell size asymmetries in the sea star embryo

**DOI:** 10.1101/2022.03.05.483134

**Authors:** Vanessa Barone, Maria Byrne, Deirdre C. Lyons

**Affiliations:** Center for Marine Biotechnology and Biomedicine, University of California San Diego, La Jolla, CA, 92093, USA; School of Life and Environmental Sciences, The University of Sydney, Camperdown, NSW 2006, Australia

**Keywords:** Cell fate, cell size, echinoderm, dorsoventral axis, embryo patterning

## Abstract

Cell size asymmetries are often linked to cell fate decisions, due to cell volumes and cell fate determinants being unequally partitioned during asymmetric cell divisions. A clear example is found in the sea urchin embryo, where a characteristic and obvious unequal 4th cleavage generates micromeres, which are necessary for mesendoderm cell fate specification. Unlike sea urchin development, sea star development is generally thought to have only equal cleavage. However, subtle cell size asymmetries can be observed in sea star embryos; whether those cell size asymmetries are consistently produced during sea star development and if they are involved in cell fate decisions remains unknown. Using confocal live imaging of early embryos we quantified cell size asymmetries in 16-cell stage embryos of two sea star species, *Patiria miniata* and *Patiriella regularis*. Using photoconversion to perform lineage tracing, we find that the position of the smallest cells of *P. miniata* embryos is biased toward anterior ventral tissues. However, both blastomere dissociation and mechanical removal of one small cell do not prevent dorsoventral (DV) axis formation, suggesting that embryos compensate for the loss of those cells and asymmetric partitioning of maternal determinants is not strictly necessary for DV patterning. Finally, we show that manipulating cell size to introduce artificial cell size asymmetries is not sufficient to direct the positioning of the future DV axis in *P. miniata* embryos. Our results show that although cell size asymmetries are consistently produced during sea star early cleavage and may be predictive of the DV axis, they are not necessary to instruct DV axis formation.

## Introduction

Echinoid embryos, including sea urchin, heart urchin, pencil urchin and sand dollars, are unique among the echinoderms with respect to their visibly asymmetric cell division at the 4th cleavage: each vegetal blastomere of the 8-cell stage embryo will divide to produce one large cell - the macromere - and one very small cell - the micromere (Hörstadius and Horstadius, 1973; Lyons et al., 2012; Schroeder, 1987). This asymmetric cell division has been best studied in the sea urchin embryo, where the unequal segregation of cell volumes is due to the spindle being positioned closer to the vegetal cortex, at the site of micromere formation (Poon et al., 2019; Schroeder,1987; Voronina and Wessel, 2006). The vegetal cortex, inherited by the micromeres only, is also enriched in cell fate determinants that induce differentiation into germ cells and primary mesenchyme cells (Henson et al., 2021; Peng and Wikramanayake, 2013; Swartz and Wessel,2015). The fact that both cell volumes and cell fate determinants are asymmetrically partitioned during the 4th cell division makes the micromeres the first morphological hallmark of the antero-posterior (AP) axis, with the micromeres positioned at the future posterior side of the embryo (Cameron et al., 1987; Hörstadius and Horstadius, 1973; Ruffins and Ettensohn, 1996). While cell fate determinants responsible for AP axis formation are also localized in the vegetal region of other echinoderm embryos, such as asteroid sea stars (Maruyama and Shinoda, 1990;Swartz et al., 2021) and brittle stars (Primus, 2005), cleavage is approximately equal in those species (Arnone et al., 2015). However, more subtle cell size asymmetries have been observed in the early embryos of asteroid sea stars (Kominami, 1983), holothuroids (Holland, 1981) and crinoids (Mladenov, 1983), raising the questions of i) how consistent and stereotypical cell size asymmetries are and ii) if cell size asymmetries are relevant to the establishment of embryonic axes in echinoderm embryos other than the sea urchin. Here we answer these questions for the asteroid sea star embryo, using a combination of high-resolution live imaging, lineage tracing and micromanipulations to quantify cell size asymmetries and their role in axis formation.

Sea star embryos present holoblastic cleavage, with the first and second cleavages both aligned with the animal-vegetal axis and the third cleavage perpendicular to the first two, and dividing animal and vegetal halves of the embryo (Byrne and Barker, 1991; Dan-Sohkawa and Satoh,1978; Kominami, 1983). The animal-vegetal axis is established during oogenesis: the germinal vesicle is asymmetrically positioned in the immature oocyte, predicting both the site of polar body extrusion and the anterior side of the embryo (Kominami, 1983). Therefore, the AP axis of the sea star embryo can be identified already in the oocytes and its establishment is thought to depend on asymmetric localization of maternal determinants (Kominami, 1983; Kuraishi and Osanai, 1992;Swartz et al., 2021; Wanninger, 2015). The first morphological hallmark of DV axis formation, instead, can be detected only at 3 days post fertilization (dpf), when the archenteron joins the anterior ectoderm to form the mouth on the ventral side of the embryo (Kominami, 1983;Wanninger, 2015).

Unlike sea urchins, early sea star embryos do not have obvious asymmetric cleavages and are generally thought to have equally sized cells (Wanninger, 2015). However, to the best of our knowledge, measurements of cell volumes in the early sea star embryo have not been performed. We find more subtle, yet consistent, cell size asymmetries in 16-cell stage sea star embryos of two species, *Patiria miniata* and *Patiriella regularis*. Using lineage tracing, we show that the position of the smallest cells in *P. miniata* embryos is biased towards the anterior-ventral side of the future embryo. Using mechanical manipulations we determined that neither maternal determinants segregated in the smallest cells, nor cell size asymmetries *per se*, are required to induce embryonic axes, suggesting that although cell size asymmetries are consistently produced during sea star early cleavage they are not necessary to instruct DV axis formation.

## Results and discussion

In euechinod sea urchins the fourth cleavage is unequal and gives rise to 4 micromeres, 4 macromeres and 8 mesomeres (Hörstadius and Horstadius, 1973; Lyons et al., 2012; Schroeder,1987). In asteroid sea stars, cleavage is thought to be equal and all blastomeres at the 16-cell stage are expected to have similar size (Wanninger, 2015). However, cell size asymmetries can be observed in a proportion of sea star embryos (Fig S1, (Kominami, 1983)).

This raises the possibility that cleavage of sea star embryos is not necessarily equal: it might produce less obvious, yet consistent, cell size asymmetries, possibly involved in axis determination. To test if sea star embryos present differently sized blastomeres and at what stage unequal cleavage might occur, we used high resolution live imaging. We analyzed embryos of two asteroid sea star species (*Patiriella regularis* and *Patiria miniata*) and one euechinoid sea urchin species (*Lytechinus pictus*) for comparison. We performed high resolution live imaging of embryos expressing membrane and nuclear markers from 4-cell stage to 16-cell stage (Fig 1A, S2, Movies S1-S3). The animal pole was assigned as opposite to the site of formation of the micromeres in *L. pictus* and as the side of polar body extrusion in the sea stars. We subsequently segmented individual cells in 3D and measured cell volumes (Fig 1A,B, S2). To compare cell size asymmetries across species with embryos of different sizes, cell volumes were normalized on embryo volume (Fig 1B).

**Fig 1.**
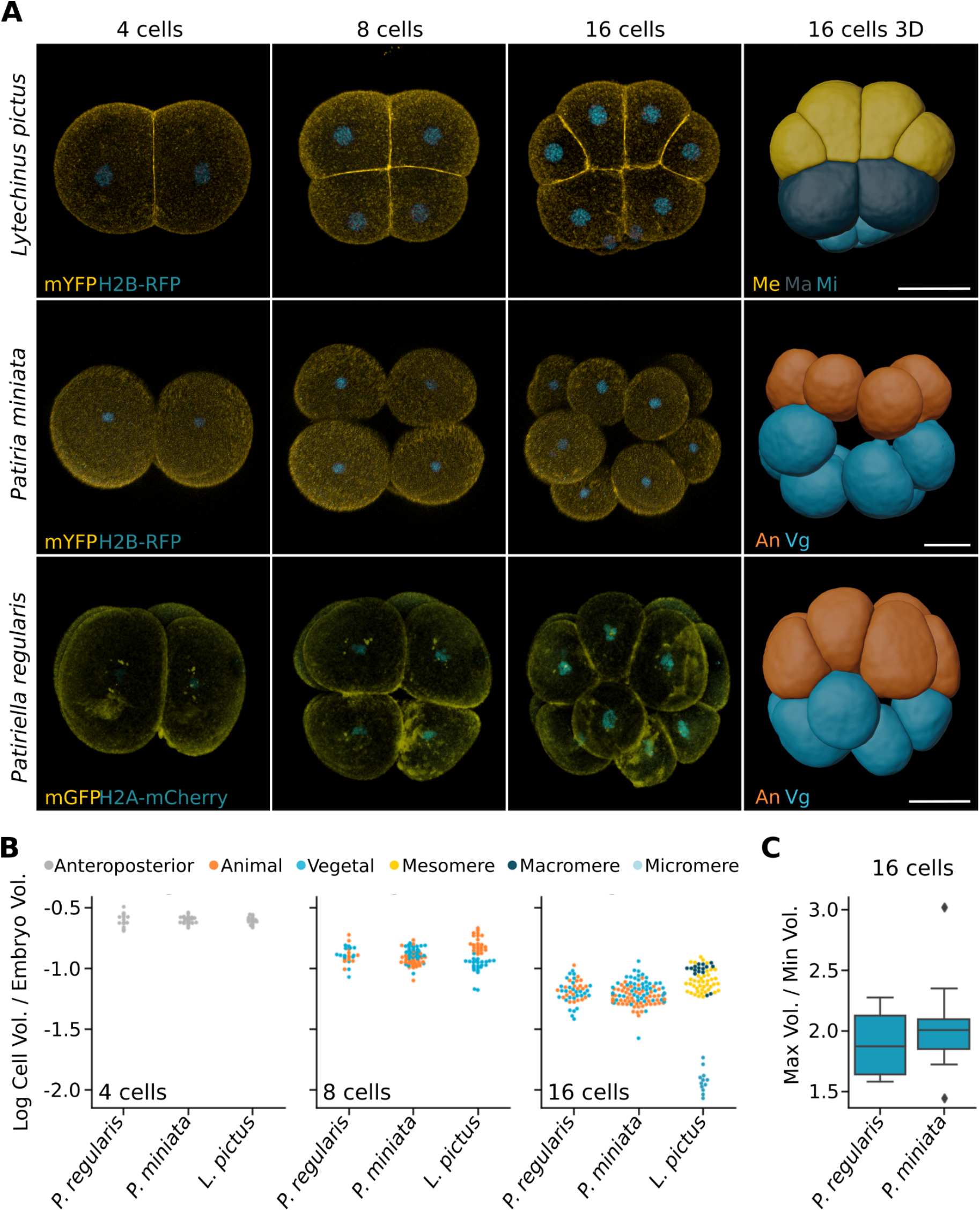
Cell size asymmetries in early sea star embryos. (**A**) Representative images of sea urchin (*Lytechinus pictus*) and sea star (*Patiria miniata, Patiriella regularis*) embryos at the 4, 8 and 16-cell stages. Embryos were injected with mRNA coding for a membrane bound fluorescent protein (mYFP, mGFP) and fluorescently tagged histone (H2B-RFP, H2B-mCherry) and subsequently imaged live on a confocal microscope. The datasets were segmented using the Fiji plugin Limeseg and individual blastomeres rendered as 3D meshes. Scale bars: 50 μm. (**B**) Volumes of individual blastomeres normalized to embryo volume, calculated as the sum of the volumes of the 4 blastomeres at the 4 cells stage. For sea star embryos, animal and vegetal poles were assigned according to the position of the polar bodies, and for sea urchin they were assigned according to the position of the micromeres. An: animal; Vg: vegetal; Me: mesomere; Ma: macromere; Mi: micromere. (**C**) Ratios of largest to smallest cells’ volume at 16-cell stage in sea star embryos. *L. pictus:* n=119 cells, 5 embryos. *P. miniata*: n=180 cells, 6 embryos; *P. regularis*: n= 91 cells, 4 embryos.

As expected, we found clear cell size asymmetries in 16-cell stage *L. pictus* embryos, with high volume variations between macromeres, mesomeres, and micromeres (Fig 1B). Interestingly, we also found variation within mesomeres, which showed a wide range of sizes (Fig 1B).

The *P. regularis* embryos have higher variation in cell size compared to both *P. miniata* and *L. pictus* embryos at the 4-cell stage and variation similar to *P. miniata* at the 8 and 16-cell stages. Interestingly, the smallest cells tend to be vegetal in *P. regularis*, while they are mostly animal in *P. miniata* (Fig 1B). Notably, the differences between larger and smaller cells in sea star embryos at 16-cell stage are comparable to the differences between macromeres and mesomeres in the sea urchin (Fig 1B), with the largest cells in *P. miniata* being on average twice the size of the smallest (max/min volume ratio: 2.05 ± 0.46; Fig 1C). Taken together, these results suggest that cell size asymmetries are consistently produced during early cleavage of the sea star embryo. In *P. miniata*, the smaller cells are more often animal, similar to observations in *P. pectinifera* (Kominami, 1983). In *P. regularis*, instead, the smaller cells are mostly vegetal. This is different from the situation in echinoid species, in which the smallest cells, the micromeres, are found on the vegetal side of all species described (Cameron et al., 1987; Hörstadius and Horstadius, 1973; Ruffins and Ettensohn, 1996; Urben et al., 1988; Wray and McClay, 1988), where they are both necessary and sufficient to induce the AP axis (Logan et al., 1999). The position of small cells between these two sea star species is, instead, variable with respect to the AP axis.

Next we asked if the position of small cells is predictive of DV axis formation in the sea star embryo. To test this hypothesis we first analyzed the relation between cleavage planes and embryonic axes in *P. miniata*. In asteroid sea stars, the embryo develops into a blastula and gastrulation consists of invagination of mesendodermal cells at the vegetal side (opposite to the polar bodies) (Kominami, 1983). During gastrula stages, the archenteron elongates within the hollow ectoderm tissues and eventually joins the anterior ectoderm to open the mouth (Byrne and Barker, 1991; Kominami, 1983). The opening of the mouth, which for *P. miniata* happens at around 72 hours post fertilization (hpf), is the first known morphological hallmark of DV axis formation: the larva is now a bipinnaria and both mouth and anus open on the ventral side (Fresques et al., 2014;Kominami, 1983; Newman, 1922). To confirm that first and third cleavages predict the AP axis of *P. miniata* larvae, we performed lineage tracing at the 2 and 8-cell stages (Fig 2). We injected one cell with a fluorescent dextran and raised the injected embryos up to the bipinnaria stage. As expected, we found that in embryos injected at the 2-cell stage, the labelled clone constituted roughly half of the larvae, including anterior ectoderm, posterior ectoderm and mesendoderm tissues (Fig 2A). We then measured the angle formed by the labelled clone with the animal-vegetal axis at the beginning of gastrulation (26 hpf) and found that the first cleavage aligns consistently with the animal-vegetal axis (Fig 2B). In embryos injected in one animal blastomere at the 8-cell stage, the labelled clone included only a portion of anterior ectoderm, while in those injected into one vegetal blastomere the labelled clone included a portion of posterior ectoderm and mesendoderm (Fig 2A,C). Next, we measured the angles formed by the clones with the DV axis at 72 hpf (Fig 3A-D) and found that both first (3A,B) and second cleavage (3C,D) are positioned randomly with respect to the DV axis.

**Fig 2.**
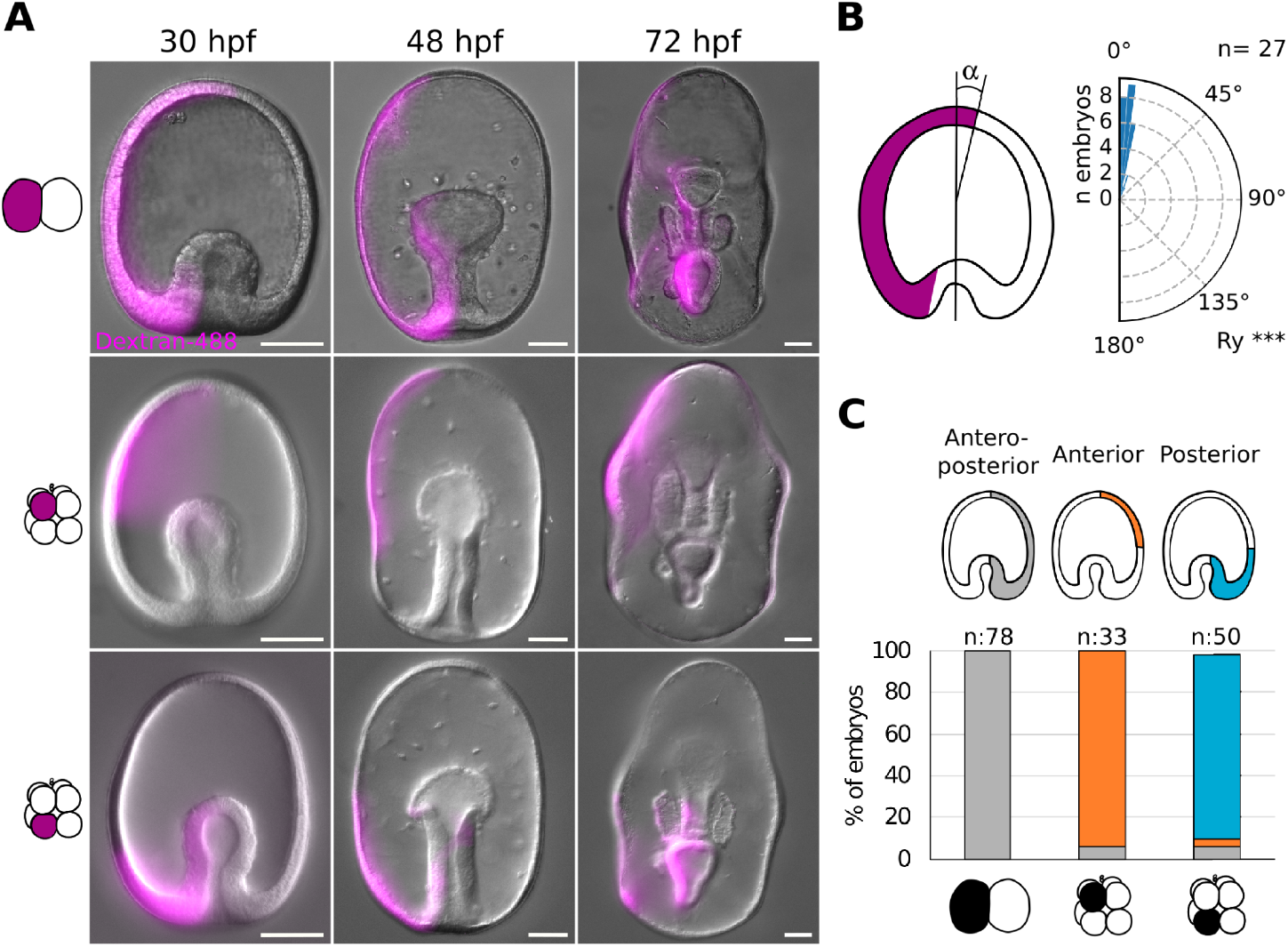
First and third cleavages predict the anteroposterior axis in *P. miniata* sea star embryo. (**A**) Representative images of *P. miniata* embryos injected with a lineage tracer at the 2 or 8-cell stage. One blastomere was injected with a mixture of Dextran-Alexa488 and mRNA coding for Histone-BFP either at the 2 or at the 8-cell stage. At the 8-cell stage, either one animal or one vegetal blastomere was injected, scored according to the position of the polar bodies. Embryos were then raised at 16C and imaged on an epifluorescence microscope at 30, 48 and 72 hpf. Scale bars: 50 μm. (**B**) Alignment of the first cleavage with the animal-vegetal axis. Embryos injected at the 2-cell stage were stained with cell mask orange and imaged in toto on a confocal microscope. The images were rendered in 3D and the angle formed between the clone and the animal-vegetal axis was measured. n= 27 embryos. Rayleigh test, ***: p-value < 0.001. (**C**) Quantification of the lineage tracing experiments. One blastomere of *P. miniata* embryos was injected either at the 2 or 8-cell stages, discriminating between animal and vegetal blastomeres at the 8 cell stage based on the position of the polar bodies. Embryos were raised at 16C and the position of the injected clone scored at 26 hpf. n= 161 embryos, 4 experiments.

**Fig 3.**
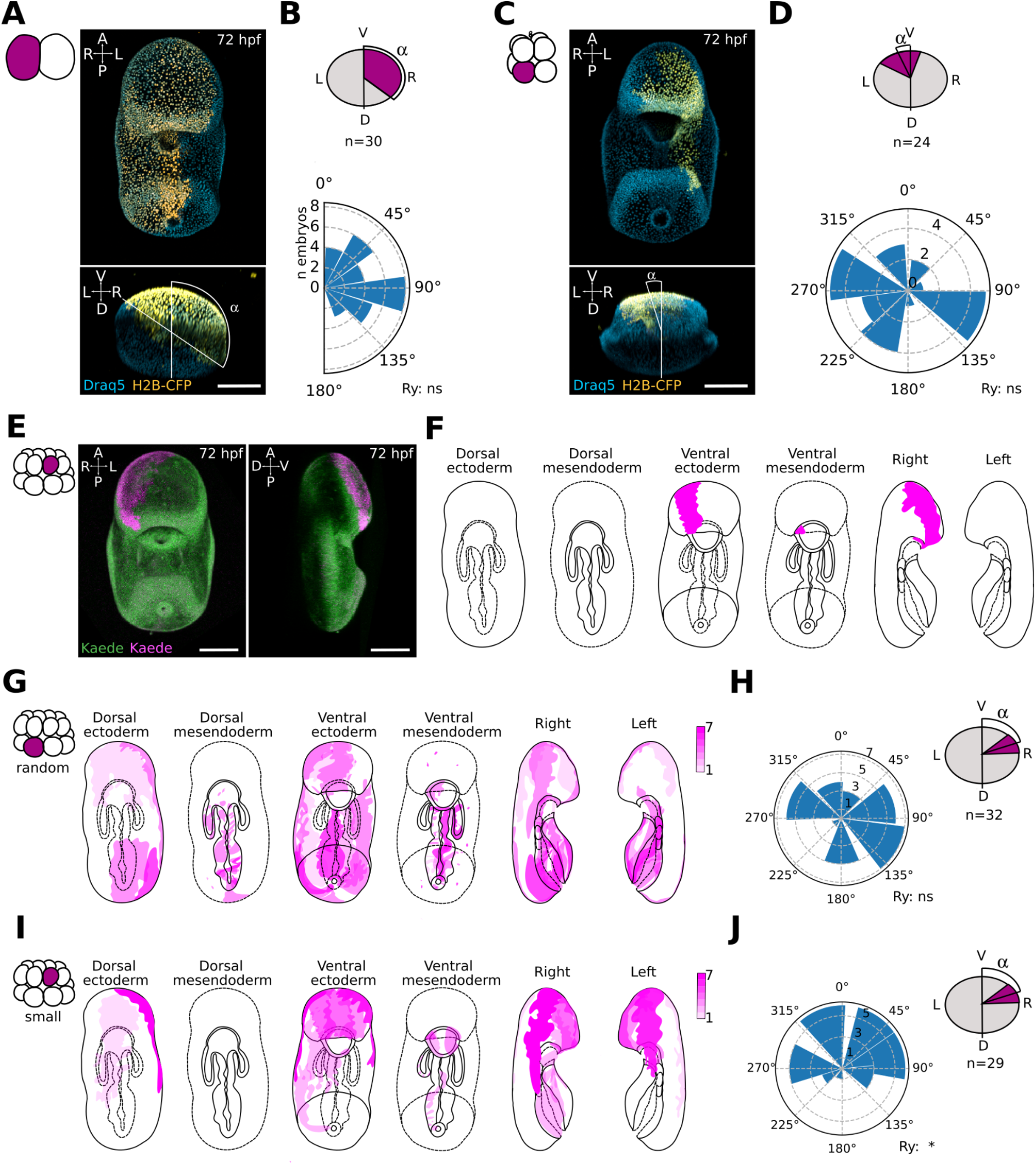
The position of smaller cells at 16-cell stage is biased toward the ventral side in *P. miniata* sea star embryos. (**A-D**) First and third cleavage in relation to the dorsoventral axis. Representative confocal image of a sea star larvae injected at the 2-cell stage (**A**) or 8-cell stage (**C**). One blastomere was injected with a mixture of Dextran-Alexa488 and mRNA coding for Histone-BFP at the 2-cell stage, raised at 16C, fixed at 72 hpf, stained with Draq5 to visualize all nuclei and imaged in toto on a confocal microscope. Quantification of the angles formed by the injected clone and the sagittal plane of the larva at 72 hpf is shown in (**B**) (2-cell; n= 30 embryos) and (**D**) (8-cell; n= 24 embryos). (**E-L**) Position of small cells in relation to the dorsoventral axis. (**E**) Representative confocal image of a sea star larva showing the clone derived by a small cell at the 16-cell stage. Oocytes were injected with mRNA coding for the photoconvertible protein Kaede (Kaede) and incubated ON at 16C. Oocytes were subsequently activated, fertilized and incubated until the 16-cell stage, when one of the 16-cell was photoconverted (Kaede) on a confocal microscope (405 nm laser). Embryos were raised at 16C for 72 hpf and then imaged live in toto on a confocal microscope. (**F**) Schematic representation of the clone shown in (**E**). (**G-J**) Quantification of the positions of clones observed after the photoconversion of a random cell (**G**) or a small cell (**I**) at 16-cell stage and the angles of the same clones formed with the sagittal plane at 72 hpf (**H**,**J**). Random cells: n= 32 embryos; Small cells: n=29 embryos. Rayleigh test, *: p-value < 0.05; ns: not significant. Scale bars: 100 μm.

Taken together, these results show that the first cleavage is aligned with the animal-vegetal axis but is not predictive of the future larval DV axis. The third cleavage separates the embryo into animal and vegetal halves, with animal cells giving rise to anterior ectoderm and vegetal cells giving rise to posterior ectoderm and mesendoderm. This confirms what was shown for *P. pectinifera* (Kominami, 1983) and our preliminary observations in *P. regularis* (Fig S3).

To test if the cell size asymmetries arising during cleavage stages are predictive of the DV axis in *P. miniata*, we turned to an imaging approach. The appearance of small cells is most obvious at the 16-cell stage in this embryo: given the difficulty of faithfully injecting those smaller cells we used photoconversion to label them. We raised embryos expressing the photoconvertible protein Kaede to the 16-cell stage and then used a UV laser on a confocal microscope to photoconvert either a random cell (control) or one small cell (Fig S4). We then raised the photoconverted embryos to the bipinnaria stage and assessed the position of the photoconverted clones (Fig 3E,F, S5, S6). We found that in larvae where a random cell had been photoconverted, the labelled clones were positioned randomly with respect to both the AP and DV axes (Fig 3G,H S5). In larvae where one small cell had been photoconverted the position of labelled clones was biased toward the ventral half of the anterior ectoderm (Fig 3I,L, S6).

Taken together, these results show that the smallest cells of *P. miniata* embryos at the 16-cell stage are more likely to arise in the animal side of the embryo, where the future ventral side will be specified. However, the association between the position of the smallest cell and the DV axis is broad, with clones deriving from the smallest cells found across the entire ventral half on the 3dpf larvae, instead of exclusively at the site of mouth opening.

Next we sought to understand if and how smaller cells may influence DV axis formation, as cell size asymmetries are often linked to cell differentiation due to asymmetric partitioning of both maternal determinants and cytoplasmic volumes during cell division (reviewed by (Sunchu and Cabernard, 2020).

To test if asymmetric partitioning of maternal determinants is necessary to DV axis determination in *P. miniata*, we performed early embryo dissociations. We isolated blastomeres at the 2, 4 and 8-cell stage and raised them to the bipinnaria stage (Fig 4, S7). As expected, control, whole embryos reached the bipinnaria stage at 72 hpf (Fig 4A,G). Blastomeres isolated at the 2-cell stage formed half-sized larvae and reached the bipinnaria stage at 72 hpf in 79.6% of cases and by 120 hpf in 84.6% of cases (Fig 4B,E,G). Blastomeres isolated at the 4 cells stage formed smaller larvae and reached the bipinnaria stage at 72 hpf in 25% of cases and by 120 hpf in 86.9% of cases (Fig 4B,E,G). This suggests that in most cases all 4 blastomeres have the potential to form the DV axis.

**Fig 4.**
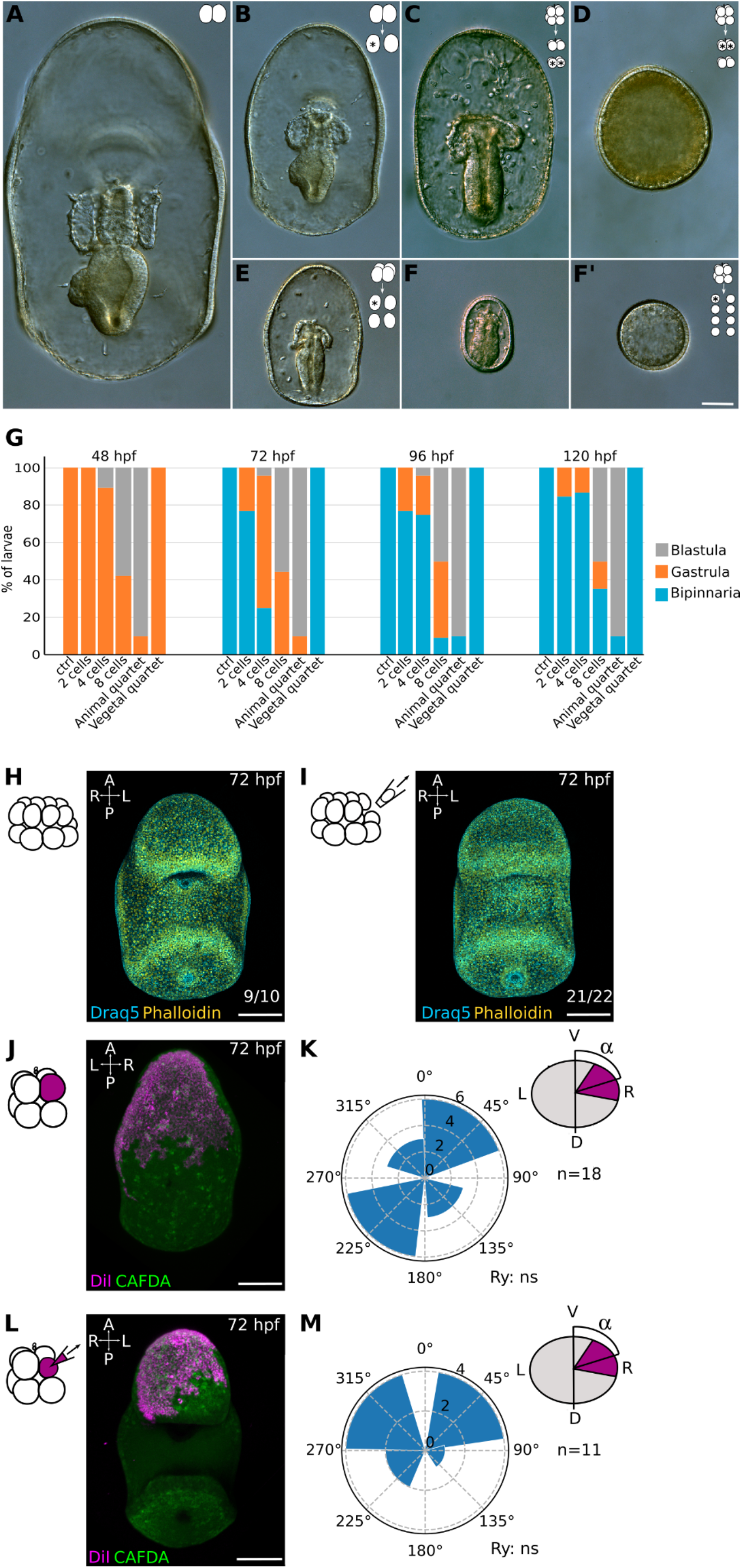
Cell size asymmetries are not necessary to establish the dorsoventral axis in *P. miniata* embryos. Representative DIC images of 72 hpf larvae generated by blastomeres of dissociated embryos. (A) CTRL embryos (not manipulated). (**B, E, F**) Larvae generated by Individual blastomeres of embryos dissociated at the 2 (**B**), 4 (**E**) or 8 (**F, F’**) cells stages. (**C,D**) Larvae generated by vegetal (**C**) or animal (D) quartets of blastomeres isolated at the 8-cell stage. This corresponds to the vegetal and animal halves of an embryo. Vegetal or animal identity was established according to the position of the polar bodies. (**G**) Phenotypes of larvae formed by isolated blastomeres at between 48 and 120 hpf. Blastula: no invagination; Gastrula: evident gut; Bipinnaria: mouth formed. n= 94 isolated blastomeres; 35 embryos; 2 experiments. (**H-I**) Removal of one small cell at the 16 cell stage. Zygotes were denuded of their fertilization envelope and raised at 16C until the 16-cell stage. No cell (**H**) or one small cell (**I**) was removed by micropipette aspiration. Embryos were raised at 16C to 72 hpf, fixed, stained with Draq5 and Phalloidin and imaged on a confocal microscope. Phenotype quantification shown on bottom right. n=33 embryos, 2 experiments. (**J-M**) Cell size reduction. Representative confocal images of CTRL (**J**) or manipulated (**L**) larvae. One blastomere at the 8-cell stage was injected with Dil and immediately reduced in size by micropipette aspiration of cytoplasm. CTRL embryos were injected but not aspirated. Quantification of the angles formed by the injected clone and the sagittal plane of the larva at 72 hpf is shown in (**K**) (CTRL; n=18 embryos) and (**M**) (Reduced; n= 11 embryos). Rayleigh test, ns: not significant. Scale bars: 50 μm.

To test if maternal determinants for DV axis formation are inherited by the animal cells at the 8-cell stage, we split embryos at the 8-cell stage into animal and vegetal quartets; we found that all the vegetal quartets developed into half-sized bipinnaria by 72 hpf (Fig 4C,G), while animal quartets failed to form mesendoderm tissues and remained blastulae for 5 days (Fig 4D,G). This suggests that the vegetal blastomeres retain the potency to establish a DV axis, even in the absence of animal cells. To test if all vegetal blastomeres are capable of establishing a DV axis, we isolated individual blastomeres at the 8-cell stage (Fig 4F,F’). We found that 44.4% of the isolated blastomeres formed small gastrulae at 72 hpf (Fig 4F) and 55.5% formed small blastulae (Fig 4F’), indicating that isolated vegetal blastomeres can form mesendoderm tissues but fail to open a mouth by 72 hpf. However, 70.5% of those gastrulating mini-larvae reached the bipinnaria stage by 120 hpf, when the experiment was terminated (Fig 4G, S7).

Taken together these results suggest that the vegetal portion of the embryo is necessary and sufficient for gut formation and establishment of the DV axis, although vegetal blastomeres isolated at the 8-cell stage establish a DV axis with considerable delay. It is worth noting that these results contradict Dan-Sohkawa and Satoh (Dan-Sohkawa and Satoh, 1978), who showed all blastomeres isolated at the 8 cell stage form gastrulae in *P. pectinifera*, but in agreement with Maruyama and Shinoda (Maruyama and Shinoda, 1990) who showed only the vegetal blastomeres form gastrulae in that same species. Interestingly, Maruyama and Shinoda find that all isolated blastomeres at the 2 and 4 cells stage, and all of the vegetal blastomeres isolated at the 8-cell stage form bipinnaria, although they do not discuss the timing of these events. Our results also align with recent experiments showing that the vegetal-most portion of cytoplasm in the oocyte is necessary for gut induction in *P. miniata* (Swartz et al., 2021). Therefore, *P. miniata* is similar to most other echinoderms analyzed so far in that differential allocation of maternal determinants is involved in the determination of the AP axis, but not necessary for the determination of the DV axis.

Zygotic cell fate determinants necessary for DV axis formation might accumulate in the small cells of *P. miniata* embryos and it is possible that cell fate determinant asymmetries are re-established after blastomeres dissociations. To test if zygotic determinants enriched in the small cells are necessary for DV axis formation, we used micropipette aspiration to remove one small cell from 16-cell stage *P. miniata* embryos (Fig 4H-I). We found that perturbed embryos opened a mouth at 72 hpf, similar to CTRL non-perturbed embryos (Fig 4H-I). Taken together these results suggest that enrichment of cell fate determinants in the small cells plays a negligible role in the determination of the DV axis in *P. miniata*.

Next we sought to test if the relation between smaller cells and DV axis formation in *P. miniata* may be due to cell size alone. If that were true, even after the smallest cells of an embryo were removed, either by aspiration or by dissociation, cell size asymmetries would still be in place, and the next smallest cell might instruct the position of the DV axis. To test this possibility, we artificially created a population of small cells on one side of the embryo, by removing cytoplasm from one of the animal blastomeres at the 8-cell stage by micropipette aspiration until that blastomere was the smallest in the embryo (Fig 4J-M). We then scored the position of the clone formed by the progeny of that miniaturized blastomere at 72 hpf. We found that miniaturized clones were randomly positioned with respect to the DV axis (Fig 4J-K), similar to CTRL embryos, in which one animal blastomere at the 8-cell stage was injected but not manipulated (Fig 4L-M). Taken together these results indicate that manipulating cell size to introduce artificial cell size asymmetries is not sufficient to direct the positioning of the future DV axis in *P. miniata* embryos.

While best-studied in echinoids, cell size asymmetries of varying degrees have been reported in embryos of most classes of echinoderms, including asteroids, holothuroids, and crinoids (Holland,1981; Kominami, 1983; Mladenov, 1983). In the sea urchin embryo, a specific protein, Activator of G-Protein Signaling, drives the asymmetric cell division that generates micromeres and macromeres, due to an evolutionary novel motif that recruits mitotic spindles to the vegetal cortex (Poon et al., 2019). That motif is absent in other echinoderm species and it is sufficient to induce asymmetric cleavage in the sea star embryo (Poon et al., 2019). The generation of micromeres in echinoids is therefore a highly controlled event that has evolved due to modifications in the sequence of key regulatory proteins.

In contrast, it is generally thought that cell size asymmetries in echinoderms other than echinoids arise randomly, possibly due to the accumulation of small asymmetries across several cleavages, and are not linked to cell fate decisions, nor to axes determination. These hypotheses, however, had never been tested. Here we provide evidence that cell size asymmetries arise consistently during early development of asteroid sea star embryos. The position of smaller cells is non-random, is species-specific and may be biased toward anterior ventral tissues in *P. miniata*. However, removal of the smaller cells does not affect development: sea star embryos can compensate for the loss of those cells, showing that asymmetric deposition of cell fate determinants in those cells is not strictly necessary for DV patterning. Moreover, the position of the DV axis is not strongly biased by experimental reduction of cell size, indicating that cell size asymmetries alone are not sufficient to instruct the positioning of embryonic axes. This situation may represent an evolutionary “in-between”, with cell size asymmetries arising consistently, but not necessarily being linked to cell fate decisions. It will be interesting to expand the analysis of cell asymmetries to other echinoderm species, with the potential of identifying other evolutionary states, ranging from completely randomly produced cell size asymmetries to other examples, more similar to the sea urchin, where differences in cell sizes are linked to cell fate decisions.

## Materials and methods

### Animal husbandry

Adult *Lytechinus pictus* were collected at La Jolla, CA, and held in free flowing seawater aquaria at a temperature of 16C. Spawning was induced by injection of 0.5M KCl, as previously described (Nesbit and Hamdoun, 2020). Adult *Patiria miniata* were purchased from Monterey Abalone Company (Monterey, CA) or South Coast Bio-Marine LLC (San Pedro, CA) and held in free flowing seawater aquaria at a temperature of 12-16C. Adult *Patiriella regularis* were collected off the coast of Tasmania (Australia) and held in aquaria at a temperature of 20C. Sea star gametes were obtained as previously described (Hodin et al., 2019). Briefly, ovaries and spermogonia were dissected via a small incision on the ventral side of adults. Sperm was stored undiluted at 4C while ovaries were fragmented to release oocytes in FSW. Maturation of released oocytes was induced by incubating for 1h at 16C in 3 μM 1-Methyladenine (Fisher Scientific, 5142-22-3). All embryos were raised in 0.22 μm filtered sea water (FSW) with the addition of 0.6 μg/ml Penicillin G sodium salt (Millipore Sigma, P3032) and 2 μg/ml Streptomycin sulfate salt (Millipore Sigma, S1277).

### mRNA, Dextran and Dil Injections

mRNAs were synthesized with the mMessage mMachine SP6 Transcription Kit (Invitrogen, AM1340). To label entire embryos, *Lytechinus pictus* were injected at the 1 cell stage with a mix of mRNAs coding for membrane bound YFP and Histone-2B-RFP (mYFP, 50 ng/μl; H2BRFP 400 ng/μl). *Patiria miniata* and *Patiriella regularis* immature oocytes were injected with mRNAs. mRNAs were injected, alone or in combination, to label membranes (mYFP or mGFP, 400 ng/μl), nuclei (H2B-RFP, 400 ng/μl; H2A-mCherry, 400 ng/μl) or cytoplasm (Kaede, 400 ng/μl). Injected oocytes were incubated at 16C overnight, activated and fertilized. To label individual blastomeres at the 2-, 4- or 8-cells stages, one blastomere of Patiria miniata and Patiriella regularis embryos was injected with either mRNAs coding for H2B-CFP (100 ng/μl), H2A-mCherry (400 ng/μl), Dextran Oregon Green 488 (1 mg/ml; Invitrogen, D7171) or Dil (Life Technologies, D282).

### Whole embryo staining

To label whole embryos for live imaging, embryos at 48 or 72 hpf were incubated with either CMFDA (1:2000, Invitrogen, C2925) or Cell Mask Green (1:10000, Invitrogen, C37608) for 30 min at RT. To label actin and nuclei, embryos at 24, 48 or 72 hpf were fixed in 2% Paraformaldehyde (PFA) at 4C overnight (ON) and stained with Alexa Fluor™ 488 Phalloidin (Invitrogen, A12379), DRAQ5 (Invitrogen, 65-0880-92) or DAPI (Invitrogen, D1306).

### Early embryo cell volumes

*Lytechinus pictus* embryos were injected with mRNAs coding for membrane and nuclear markers on a glass bottom dish (MatTek, P35G-1.5-14-C) coated with protamine, incubated at 16C until the 2-cell stage and then imaged on an inverted Leica Sp8 confocal microscope (20X objective, NA 0.7, 16C controlled temperature) until the 16-cell stage. *Patiria miniata* and *Patiriella regularis* embryos expressing membrane and nuclear markers were mounted on a glass bottom dish. No medium was used to immobilize the embryos: the glass bottom part of the dish was covered with a coverslip and sealed with vaseline. This creates a 1 mm deep chamber in which capillarity prevents the embryos from moving, until they develop cilia. Additional FSW was added in the dish, to help with temperature control. Embryos can be cultured in such chambers without apparent defects up to 3 dpf, when they would start feeding. Embryos were incubated until the 2 cell stage and then images on an inverted Leica Sp8 confocal microscope (20X objective, NA 0.7, 16C controlled temperature for *Patiria miniata*) or Zeiss LSM 800 confocal microscope (20X Objective, NA 0.8, 20C controlled temperature for *Patiriella regularis*) until the 16-cell stage. Datasets were 3D rendered using Imaris 6.4 (Bitplane), and further analysed using the Fiji plugin Limeseg (Machado et al., 2019) and custom python scripts. Briefly, 3D meshes for individual blastomeres at the 4-, 8- and 16-cells stages were computed with the Limeseg plugin, exported in .ply format and their volume was calculated in python. To compare cell sizes across species, cell volumes were normalized on embryo volumes, calculated as the sum of the volumes of the 4 cells at the 4 cells stage for each embryo analysed.

### Lineage tracing

To establish how early cleavage planes relate to the AP axis, one blastomere of 2- or 8-cells stage *Patiria miniata* embryos was injected with Dextran-488. Embryos injected at the 8-cells stage were separated into two groups immediately after injections based on the position of the injected blastomere: close to the polar body was considered animal, opposite to polar bodies was considered vegetal. Embryos were raised to 72 hpf and the position of the labelled clone was scored at 30, 48 and 72 hpf. A subset of embryos was live imaged at the same stages with an upright Zeiss Imager M2 (20X objective, NA 0.7). To establish how early cleavage planes are related to the DV axis, one blastomere of 2- or 8-cells stage *Patiria miniata* embryos were injected with mRNA coding for a nuclear marker. Embryos were raised until the 72 hpf, fixed, stained with DRAQ5 and imaged on an inverted Leica Sp8 confocal microscope (20X objective, NA 0.7). Datasets were 3D rendered with Imaris 6.4 (Bitplane) to measure the angle between the labelled clone and the sagittal plane. To determine how the position of the smaller cells in *Patiria miniata* relates to the DV axis, embryos expressing the photoconvertible protein Kaede were raised until the 16-cell stage, mounted on a glass bottom dish and imaged with an inverted Zeiss LSM 710 confocal microscope (20X objective, NA 0.8). Embryos are oriented randomly with this method and the cell that happened to be in optimal position for photoconversion, i.e. tilted so as not to photoconvert other cells above it - was was photoncoverted using the bleaching tool in the Zeiss Blue software: 405 nm laser at 1% power emission was used to scan a ROI inside the target cells for up to 60 sec. As control (CTRL), any embryo was photoconverted; as experimental condition (small cell), only embryos in which the smallest cell was in the optimal position was photoconverted. The smallest cells was identified by measuring the three major axis compared with the other cells of the embryo. Photoconverted embryos were recovered, raised until 72 hpf and imaged live on an inverted Leica Sp8 confocal microscope (20X objective, NA 0.7). Datasets were 3D rendered using Imaris 6.4 (Bitplane) to measure the angle between the photoconverted clone and the sagittal plane. Schematic representations of all clones were drawn based on the 3D renderings to identify clones that formed similar structures in multiple larvae.

### Mechanical manipulation of sea star embryos

To perform embryo dissociations, *Patiria miniata* oocytes were fertilized in 10 mM 4-Aminobenzoic acid (PABA, Sigma, A9878) in FSW and embryos were incubated in 10 mM PABA in FSW for 1 hpf. The fertilization envelope was mechanically removed by passing the embryos through a narrow glass pipette. Denuded embryos were transferred to gelatin coated dishes and raised in FSW until the 2-, 4- or 8-cells stage. Blastomeres were separated using an entomology needle and/or passing the embryos through a narrow glass pipette coated with gelatine. Individual blastomeres were raised separately and their phenotypes scored at 48, 72, 96 and 120 hpf. A subset of dissociated embryos was live imaged at the same stages on an upright Zeiss Imager M2 microscope (20X objective, NA 0.8). To remove one small cell at the 16 cell stage, denuded embryos were cultured in FSW on gelatine dishes and manipulated with glass micropipettes connected to a syringe. Two micropipettes were used, one with an opening diameter of 80 μm to orient and hold the embryo and the second with an opening diameter of 20 μm to remove one small cell by suction. The manipulated embryos were raised until the 72 hpf, fixed, stained with Phalloidin-488 and DRAQ5 and imaged on an inverted Leica Sp8 confocal microscope (20X objective, NA 0.7). To create a population of artificially small cells, one animal blastomere at the 8-cell stage was injected with Dil and part of its cytoplasm was suctioned away using a glass micropipette with an opening of about 5 μm. CTRL (injected but not reduced) and reduced embryos were raised until the 72 hpf stage, stained with either Cell Mask Green or CMFDA and live imaged on an inverted Leica Sp8 confocal microscope (20X objective, NA 0.7). To perform embryo dissociation of *P. regularis*, fertilization envelopes were removed mechanically at 1-cell stage, embryos were raised until the 8-cells stage and passed through a 60 μm nylon mesh. Individual blastomeres were raised for 48h at 20C and imaged.

### Statistical analysis

Statistical analyses of data were performed using Python scripts as indicated in the figure captions. The Rayleigh test was performed to assess bias in the positions of labelled clones following lineage tracing experiments in Figs 2, 3 and 4. No statistical method was used to predetermine sample size, the experiments were not randomized and the investigators were not blinded to allocation during experiments and outcome assessment.

## Acknowledgments

We are grateful to Joaquin Navajas Acedo, Victor Vacquier, and to all members of the Lyons, Byrne and Hamdoun groups for helpful discussions on the project and feedback during manuscript preparation.

## Data availability

All microscopy data is available upon request.

