## Supplementary Figures and Legends for "Cell size asymmetries in the sea star embryo"

**Fig S1**

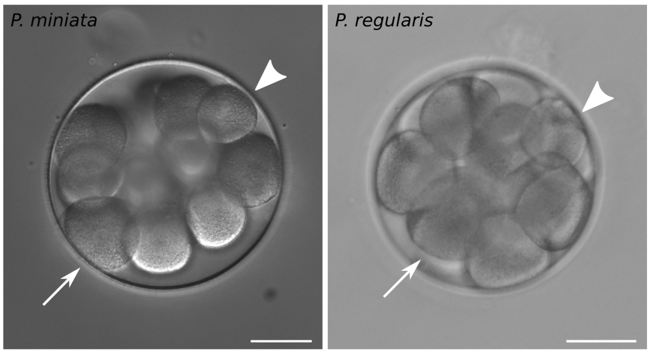


**Fig S1. Cell size asymmetries in asteroid sea star embryos.** Representative DIC images of 16-cells stage embryos of *P. miniata* (**A**) and *P. regularis* (**B**). Arrowheads point at small cells and arrows point at large cells. Scale bars: 50 μm.

**Fig S2**

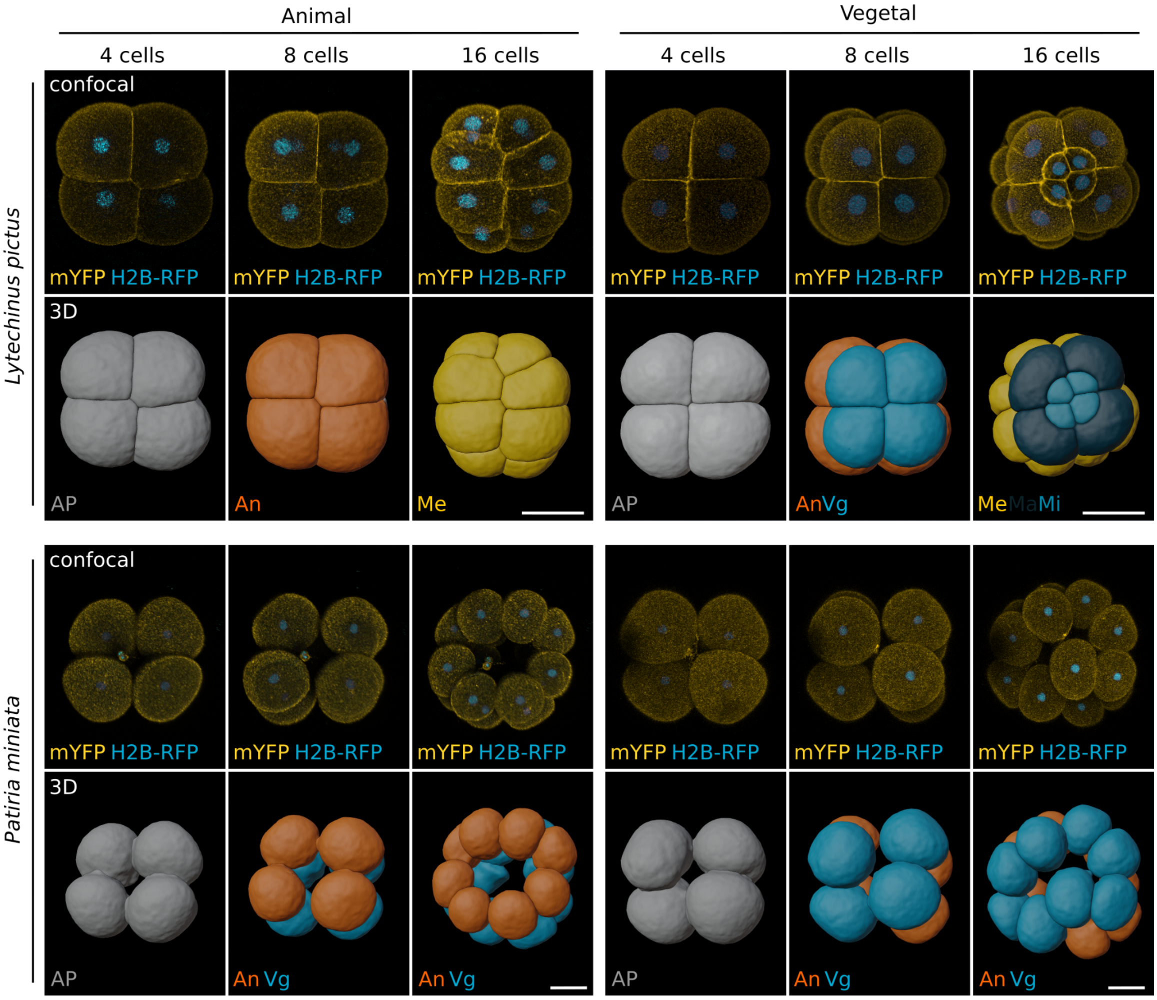


**Fig S2. 3D reconstructions of early echinoderm embryos.** Representative images of sea urchin (*Lytechinus pictus*) and sea stars (*Patiria miniata*) embryos at the 4, 8 and 16-cell stages. Embryos were injected with mRNA coding for a membrane bound YFP (mYFP) and fluorescently tagged histone (H2B-RFP) and subsequently imaged live on a confocal microscope. The datasets were segmented using the Fiji plugin Limeseg and individual blastomeres rendered as 3D meshes. Scale bars: 50 μm.

**Fig S3**

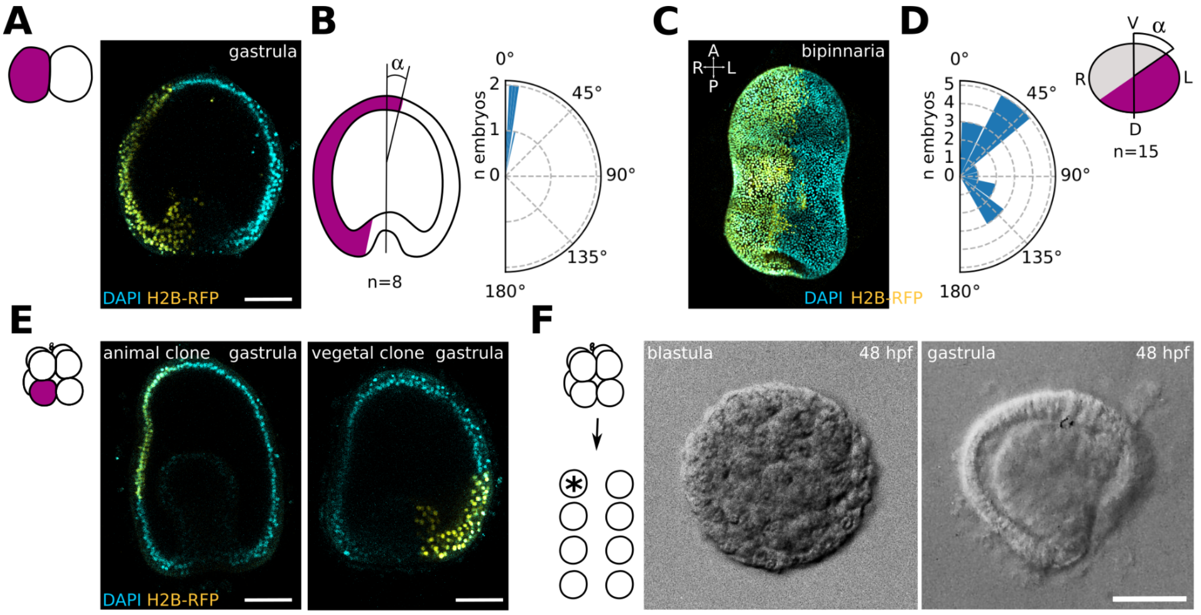


**Fig S3. First and third cleavages predict the anteroposterior axis in *P. regularis* sea star embryo.** (**A-D**) *P. regularis* embryos injected with a lineage tracer at the 2-cells stage. One blastomere was injected with mRNA coding for Histone-RFP at the 2-cells stage. Embryos were then raised at 20C, fixed, stained with Draq5 (nuclei) and imaged in toto on a confocal microscope at gastrula and bipinnaria stages. (**A**) Representative image of an injected embryo at the gastrula stage. (**B**) Alignment of the first cleavage with the animal-vegetal axis. Images of gastrula stage embryos were rendered in 3D and the angle formed between the clone formed by the injected blastomere and the animal-vegetal axis was measured. n= 8 embryos. (**C**) Representative image of an injected embryo at the bipinnaria stage. (**D**) Alignment of the first cleavage with the DV axis. Images of bipinnaria stage embryos were rendered in 3D and the angle formed between the clone formed by the injected blastomere and the sagittal plane was measured. n= 15 embryos. (**E**) Representative images of *P. regularis* embryos injected with a lineage tracer at the 8-cells stage. One blastomere was injected with mRNA coding for Histone-RFP at the 8-cells stage, embryos were then raised at 20C until gastrula stage, fixed, stained with Draq5 (nuclei) and imaged in toto on a confocal microscope. Two types of clones were observed, either forming anterior ectoderm or posterior ectoderm and mesendoderm tissues. (**F**) Representative images of *P. regularis* embryos formed by individual blastomeres separated at the 8-cells stage. Fertilization envelopes were removed mechanically at 1-cell stage, embryos were raised until the 8-cells stage and dissociated by passing them through a 60 μm nylon mesh. Individual blastomeres were raised for 48h at 20C. Two types of embryos were observed, blastulae and gastrulae. Scale bars: 50 μm.

**Fig S4**

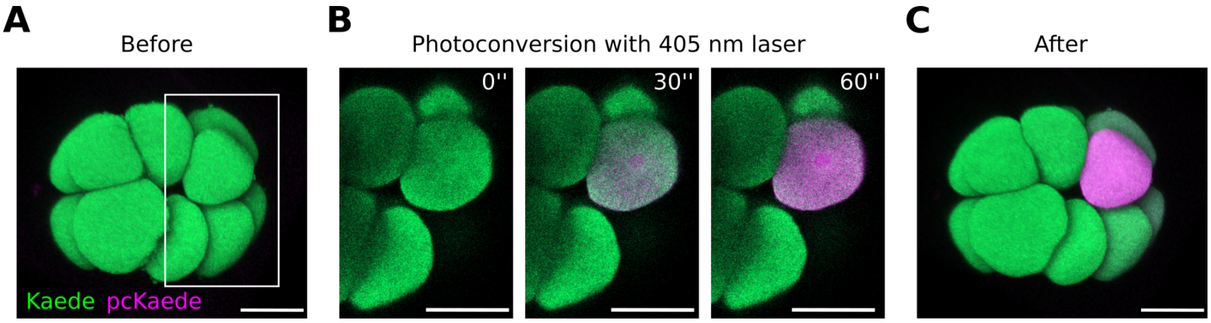


**Fig S4. Photoconversion of Kaede expressing sea star embryos.** Representative confocal images of a photoconversion experiment. *P. miniata* oocytes were injected with mRNA coding for the photoconvertible protein Kaede (Kaede) and incubated ON at 16C. Oocytes were subsequently activated, fertilized and incubated until 16-cell stage, when one of the 16-cells was photoconverted (pcKaede) on a confocal microscope (405 nm laser). (**A**) 3D rendering of a 16-cells stage embryo before photoconversion. (**B**) Close up images of the photoconverted cell after 0, 30 and 60 seconds of exposure to a 405 nm laser. (**C**) 3D rendering of the same embryo after photoconversion. Scale bars: 50 μm.

**Fig S5**

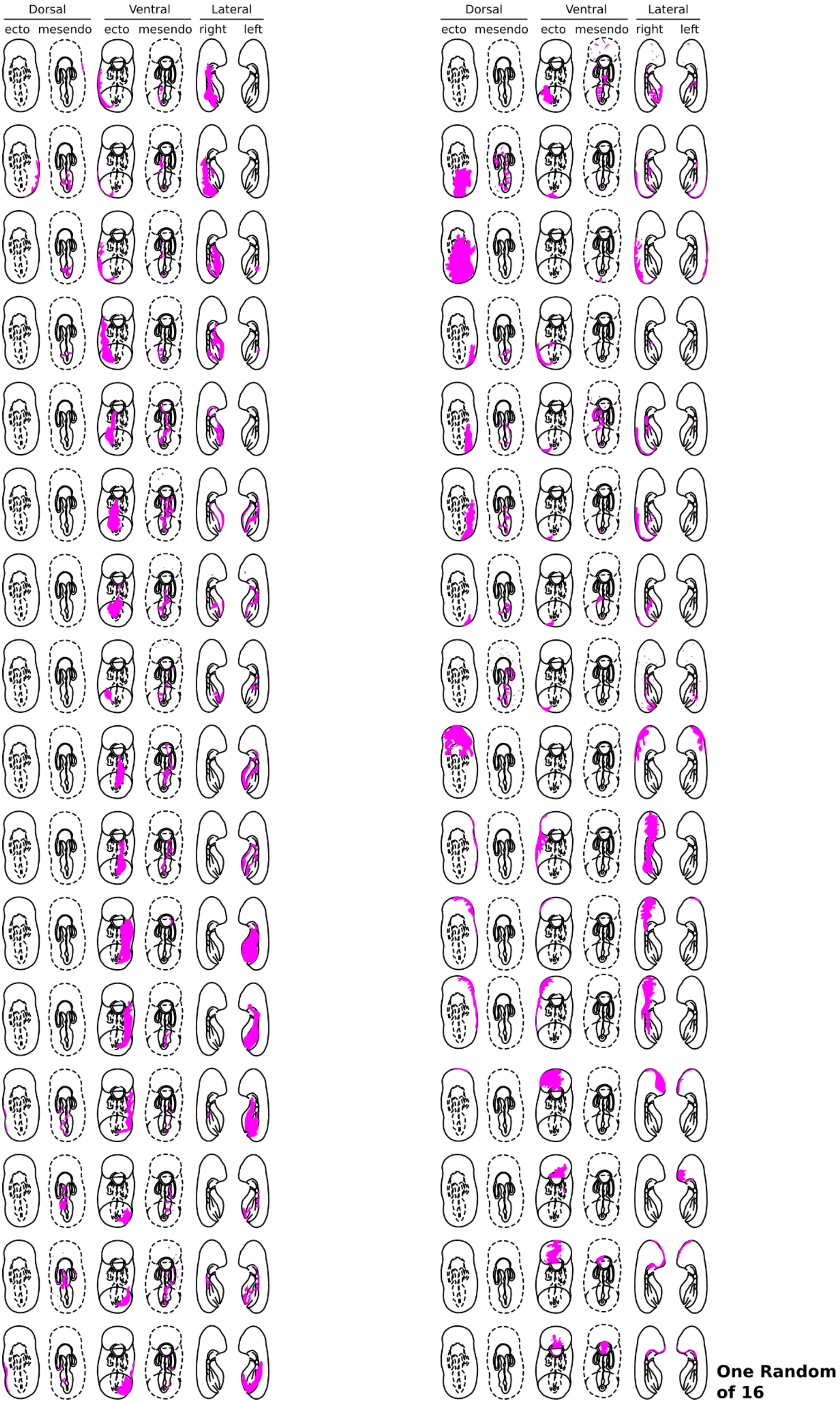


**Fig S5. *P. miniata* lineage tracing at 16-cells stage: photoconversion of one random cell.** Schematic representations of sea star larvae showing the clone derived by one cell at 16-cell stage. Oocytes were injected with mRNA coding for the photoconvertible protein Kaede (Kaede) and incubated ON at 16C. Oocytes were subsequently activated, fertilized and incubated until 16-cell stage, when one of the 16-cell was photoconverted (Kaede) on a confocal microscope (405 nm laser). Embryos were raised at 16C for 72 hpf and then imaged live in toto on a confocal microscope. Images were 3D rendered and schematic representations of the clones were drawn for easier comparison. Each clone (rows) is represented by dorsal, ventral and lateral views and distinguishing between ectodermal and mesendodermal tissues.

**Fig S6**

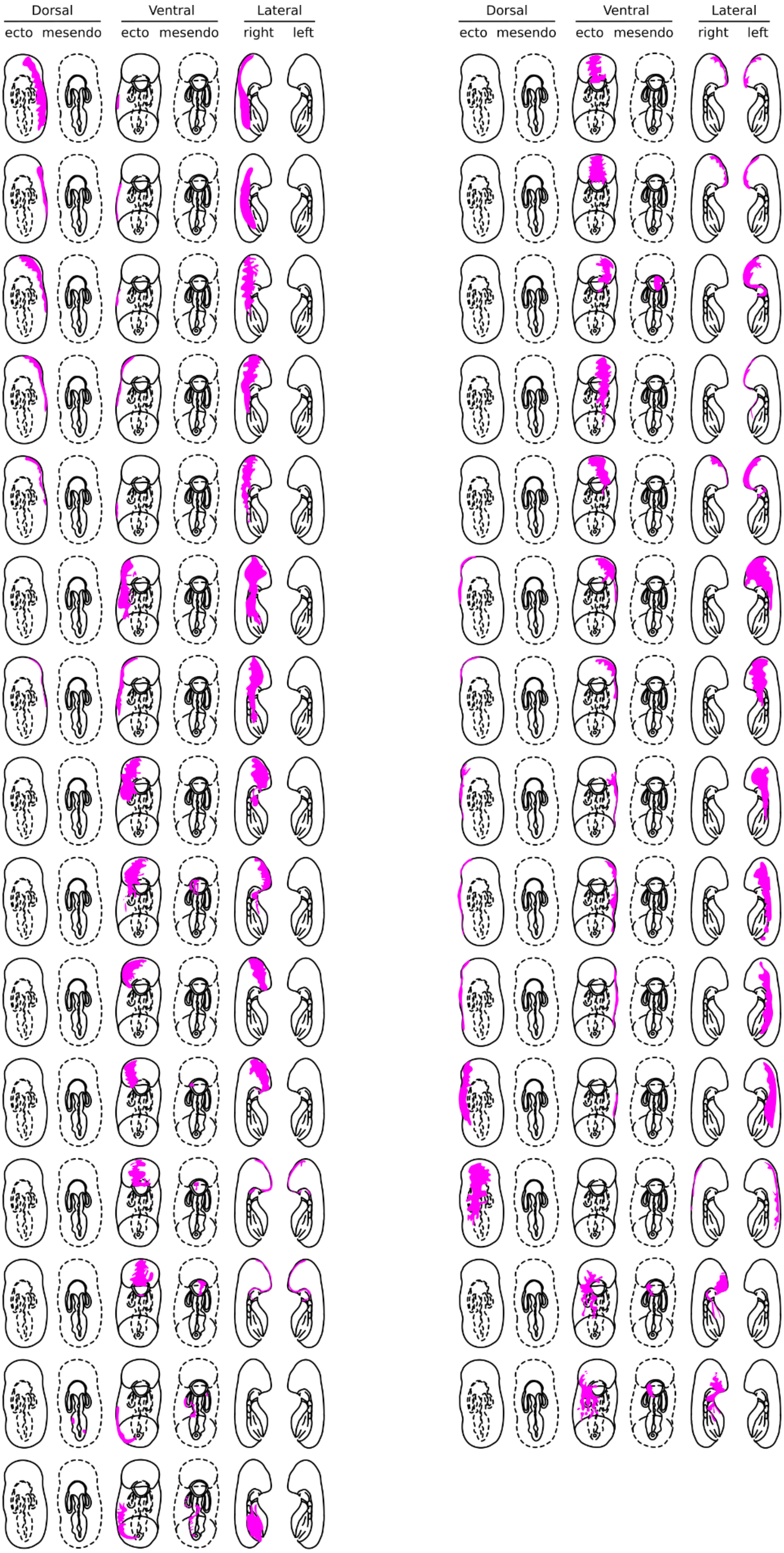


**One Small of 16**

**Fig S6. *P. miniata* lineage tracing at 16-cells stage: photoconversion of one small cell.** Schematic representations of sea star larvae showing the clone derived by one small cell at 16-cell stage. Oocytes were injected with mRNA coding for the photoconvertible protein Kaede (Kaede) and incubated ON at 16C. Oocytes were subsequently activated, fertilized and incubated until 16-cell stage, when one of the 16-cell was photoconverted (Kaede) on a confocal microscope (405 nm laser). Embryos were raised at 16C for 72 hpf and then imaged live in toto on a confocal microscope. Images were 3D rendered and schematic representations of the clones were drawn for easier comparison. Each clone (rows) is represented by dorsal, ventral and lateral views and distinguishing between ectodermal and mesendodermal tissues.

**Fig S7**

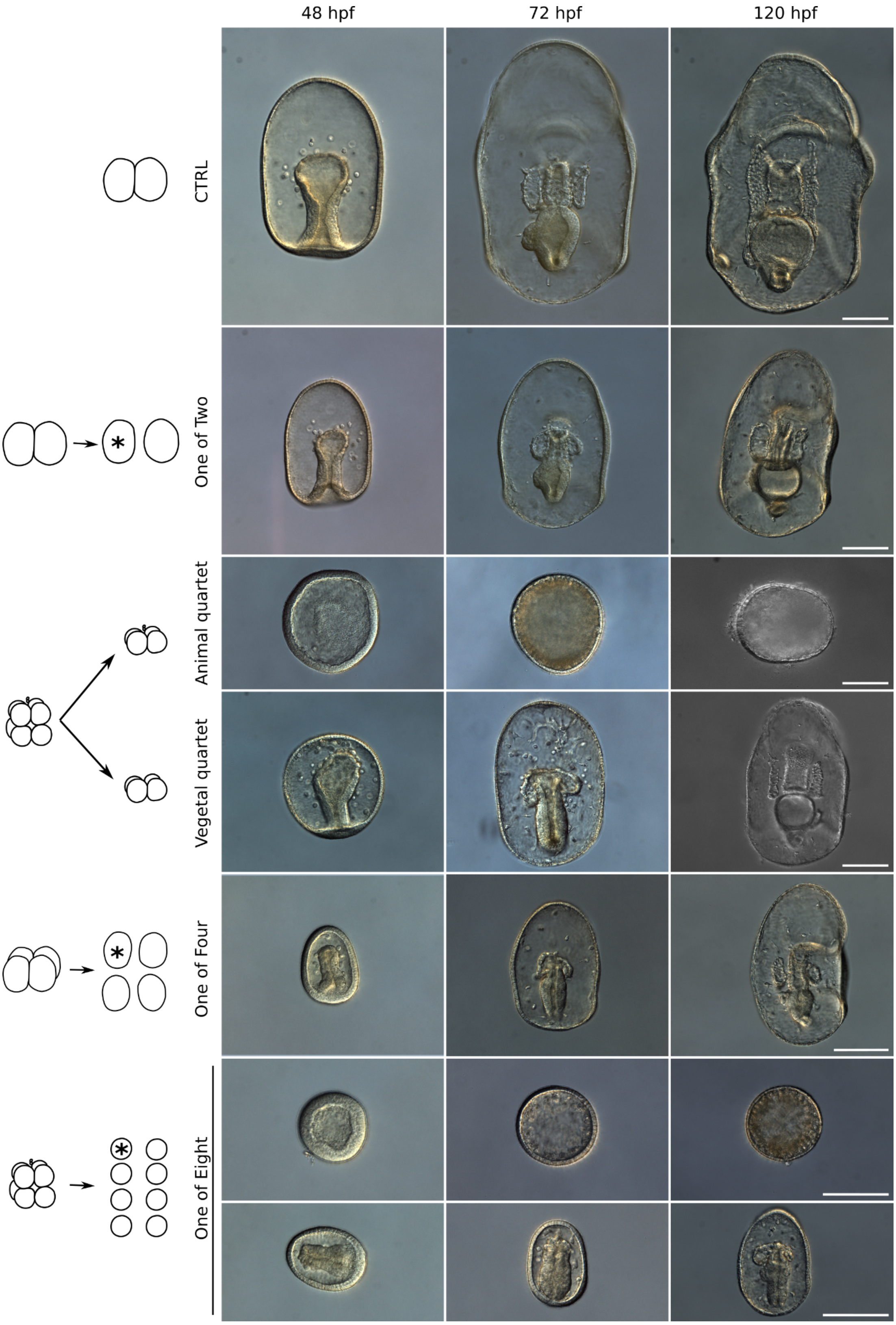


**Fig S7. *P. miniata* blastomere dissociations.** Representative DIC images of larvae generated by blastomeres of dissociated embryos. Embryos were either not manipulated (CTRL) or dissociated at the 2-, 4-, or 8-cells stage. Individual blastomeres generated by dissociation at the 2-, 4-, and 8-cells stage, and animal and vegetal quartets generated by halving embryos at the 8-cells stage were raised until 120 hpf. A subset of embryos was live imaged at 48, 72 and 120 hpf. Vegetal or animal identity was established according to the position of the polar bodies. Scale bars: 100 µm.

**Movie S1. Early *Lytechinus Pictus* embryo.** 3D rendering video of a live imaged sea urchin embryo (*Lytechinus pictus*) Embryos were injected with mRNA coding for a membrane bound fluorescent protein (mYFP, yellow) and fluorescently tagged histone (H2B-RFP, cian) and subsequently imaged live on a confocal microscope. A lateral view is shown, animal side on top.

**Movie S2. Early *Patiria miniata* embryo.** 3D rendering video of a live imaged sea star embryo (*Patiria miniata*) Oocytes were injected with mRNA coding for a membrane bound fluorescent protein (mYFP, yellow) and fluorescently tagged histone (H2B-RFP, cian), incubated ON at 16C and subsequently activated, fertilized and imaged live on a confocal microscope. A lateral view is shown, animal side on top.

**Movie S3. Early *Patiriella regularis* embryo.** 3D rendering video of a live imaged sea star embryo (*Patiriella regularis*) Oocytes were injected with mRNA coding for a membrane bound fluorescent protein (mGFP, yellow) and fluorescently tagged histone (H2A-mCherry, cian), incubated ON at 16C and subsequently activated, fertilized and imaged live on a confocal microscope. A lateral view is shown, animal side on top.

**Movie S4. Photoconverted *P. miniata* larva.** Animation showing a 3D reconstruction of a representative embryo photoconverted at the 16-cells stage. Oocytes were injected with mRNA coding for the photoconvertible protein Kaede (Kaede) and incubated ON at 16C. Oocytes were subsequently activated, fertilized and incubated until 16-cell stage, when one of the 16-cell was photoconverted (Kaede) on a confocal microscope (405 nm laser). Embryos were raised at 16C for 72 hpf and then imaged live in toto on a confocal microscope. Images were 3D rendered with Imaris (Bitplane) and animation rotating the dataset was recorded. Kaede is shown in green and photoconverted Kaede in red.
